# The application of mixed linear models for the estimation of functional effects on bovine stature based on SNP summary statistics from a whole-genome association study

**DOI:** 10.1101/2022.06.11.495744

**Authors:** Krzysztof Kotlarz, Barbara Kosinska-Selbi, Zexi Cai, Goutam Sahana, Joanna Szyda

## Abstract

Genome-Wide Association Studies (GWAS) help identify polymorphic sites or genes linked to phenotypic variance, but a few identified genes / Single Nucleotide Polymorphisms are unlikely to explain a large part of the phenotypic variability of complex traits. In this study, the focus was moved from single loci to functional units, expressed by the metabolic pathways: Kyoto Encyclopaedia of Genes and Genomes (KEGG). Consequently, this study aimed to estimate KEGG effects on stature in three Nordic dairy cattle breeds using SNPs effects from GWAS as the dependent variable. The SNPs were annotated to genes, then the genes to KEGG pathways. The effects of KEGG were estimated separately for each breed using a mixed linear model incorporating the similarity between pathways expressed by common genes. The KEGG pathway D-amino acid metabolism (map00473) was estimated as significant on stature in two of the analysed breeds and revealed a borderline significance in the third breed. Interestingly, biological evidence exists that described the importance of D-amino acids for growth in experimental organisms as well as in cattle.

## Introduction

Genome-Wide Association Studies (GWAS) are very useful for the identification of polymorphic sites, typically Single Nucleotide Polymorphisms (SNPs), or sometimes genes associated with a phenotypic variation or with a disease. Nowadays, the common availability of SNPs obtained based on whole-genome sequencing allows for a very good resolution of the estimation of those associations. However, in the context of phenotypes undergoing a complex mode of inheritance, it is not expected that a few genes / SNPs suffice to explain the variability on a phenotypic level. As a consequence, we often manage to identify loci with a very high effect on the phenotypic variation, but still, a predominant proportion of this variation remains unexplained (Manolio et al. 2009), since it is often due to a combined effect of many loci, each with a moderate or small impact. Therefore, in our study, we moved the focus from individual locus to functional units, here expressed by the metabolic pathways defined by the Kyoto Encyclopaedia of Genes and Genomes (KEGG) database. This approach allows us to better understand the physiological mechanisms underlying complex phenotypes. For this purpose, we used SNP summary statistics originating from the GWAS conducted for stature and based on whole-genome sequence data of three Nordic dairy cattle breeds.

## Results

The effects of 179 KEGG pathways were estimated based on the effects of selected SNPs from a whole-genome sequence-based GWAS of Bouwman et al. (2018), separately for three Nordic cattle breeds - Danish Holstein (DH with 366,877 SNPs), Danish Red Dairy Cattle (DR with 299,723 SNPs), and Finnish Red Dairy Cattle (FR with 396,224 SNPs) (Figure 1). In two breeds, the same pathway - D-amino acid metabolism (map00473) revealed a significant effect on stature with moderate P-values of 0.035 in FR and 0.049 in DH. In DR it also reached a borderline significance of 0.133. Depending on the breed, the effect of map00473 was estimated based on 78 SNPs in DH and FR, and 76 SNPs in DR (Figure 2, Supplemental Data S1). The differences in SNP counts resulted from the fact that the input SNP panel in Bouwman et al. (2018) was pre-processed separately for each breed, which resulted in breed-specific SNP exclusion.

**Figure 1.**
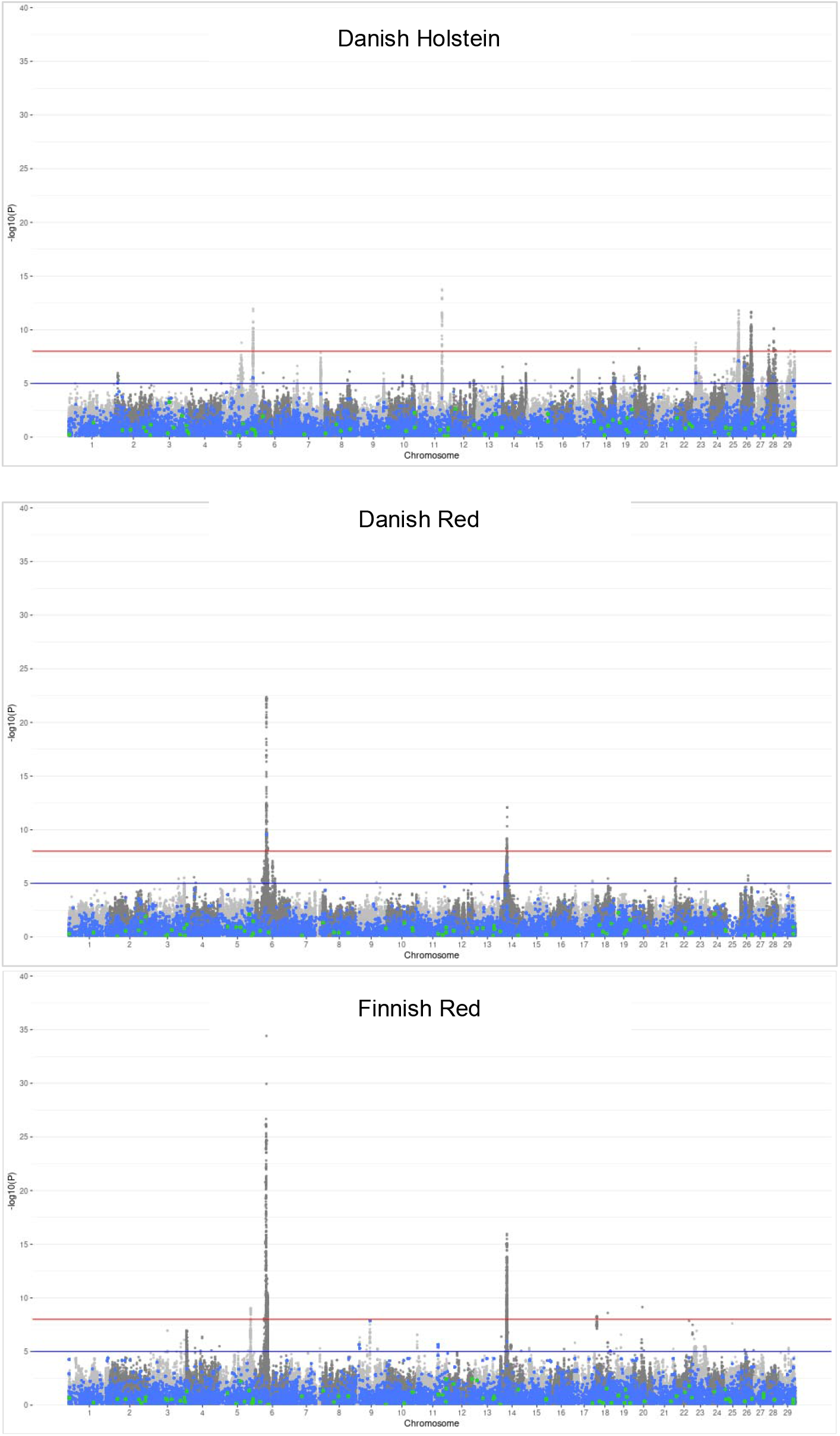
SNP significance from the whole-genome sequencing study of Bouwman et al. (2018). Blue dots correspond to genic SNPs used for the estimation of KEGG pathway effects in model (1), green dots correspond to SNPs marking genes constituting the map00473 pathway, and gray SNPs are the remainder.

**Figure 2.**
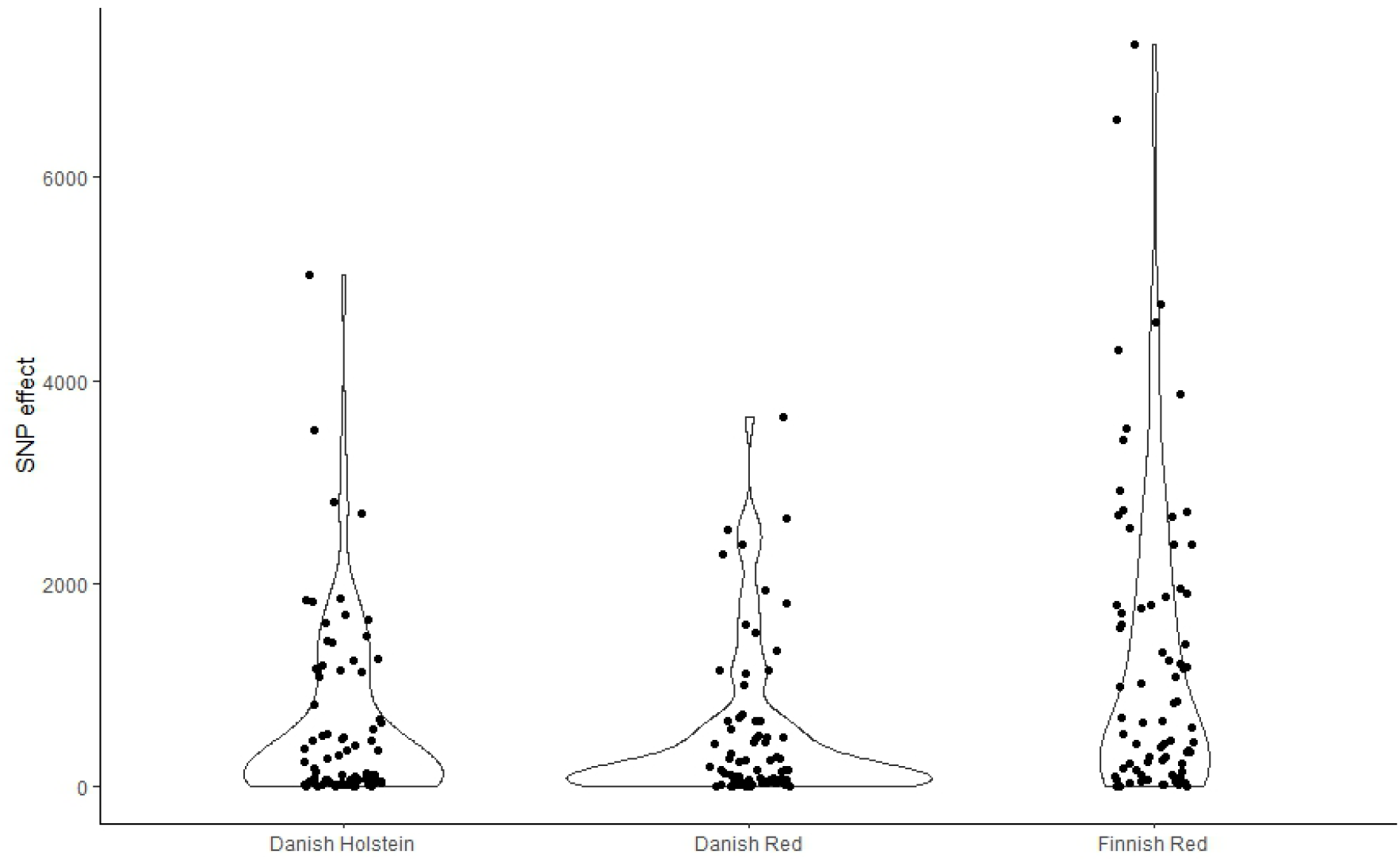
Estimated effects of SNPs marking genes from the map00473 pathway. Additionally, the pathway responsible for the metabolism of terpenoids and polyketides (map01059) was significant (P=0.041) in DH, while the synthesis and degradation of ketone bodies pathway (map00072) and the pathway of biosynthesis of various plant secondary metabolites (map00999) were significant in DR with P=0.037 and P=0.047 respectively.

## Discussion

While interpreting KEGG pathway effects two scenarios emerge. On the one hand, the overall high effect of a pathway may be driven by a high effect of a single gene that is this pathway’s component – a situation that could have been detected in a conventional genome-wide association study (GWAS). On the other hand, the high pathway effect may be due to the combined effects of many genes constituting this pathway – a situation that may easily be missed in GWAS due to the small or moderate effects of particular genes from the pathway.

In the case of our data – none of the genes harbouring the most significant SNPs in GWAS performed by Bouwman et al. (2018) was the component of the D-amino acid metabolism pathway, which therefore leads to the conclusion that the whole pathway is a significant component of the genetic determination of stature. Biologically, an outstanding pattern of our study was that the pathway associated with the metabolism of D-amino acids in all three breeds is significant for two breeds and on the border of claimed significance in the third breed. Although D-amino acids do not occur in naturally translated proteins, the link between D-amino acids metabolism and growth has long been recognised. Experimentally, a supplementation of mice with D-amino acids resulted in increased weight observed with the increased concentration of D-Phenylalanine and D-Tryptophan in the diet (Friedman and Levin, 2012). Moreover, D’Aniello (2007) reported that, in the pituitary gland, D-aspartic acid stimulates the secretion of the growth hormone in rats. In cattle, a supplementation of food with synthetic amino acids is a very common practice with commercial diet supplements containing a mixture of naturally occurring L-versions as well as not naturally occurring D-version. Campbell et al. (1996) observed that D-amino acids are somewhat less efficiently metabolised than their naturally occurring synonyms. Since methionine is often the first limiting amino acid for growth in cattle (Richardson and Hatfield, 1978) individuals that possess a more efficient mechanism of D-amino acid metabolism are expected to grow better which may result in higher stature in adults.

Another metabolic pathway demonstrating potential importance on stature is the synthesis and degradation of the ketone bodies pathway (map00072) that was significant in DR. It has been demonstrated that ketone bodies metabolism is related to growth on the whole organism (mainly through the *SLC16A6* gene as reported by Kichaev et al. (2019) and Karanth et al. (2019)) as well as on the single-cell level (Kolb et al. 2021). Moreover, although the other significant pathway of biosynthesis of various plant secondary metabolites does not directly relate to animal metabolism, it can be hypothesised that genes playing a role in the biochemical processing of metabolites originating from plants lead to higher feed efficiency in cattle and furthermore influence animals growth, but experimental evidence is lacking. Still, our results demonstrate that considering higher-order components of biological systems, such as metabolic pathways, provides a valuable insight into the basis of the variation of complex phenotypes, that may be missed by conventional GWAS analyses and should be used as an enhancement thereof.

## Materials and Methods

### Material

The analysed data comprised SNP summary statistics from GWAS performed on 5,062 Danish Holstein bulls, 924 Danish Red Dairy Cattle bulls, and 2,122 Finnish Red Dairy Cattle bulls (Bouwman et al. 2018). The association was calculated for 25.4 million variants imputed with Minimac2 (Fuchsberger et al. 2015) from 630,000 SNPs using the 1000 Bull Genomes reference population from Run4, consisting of 1,147 individuals. SNP additive effects were estimated for deregressed EBVs serving as pseudophenotypes, separately for each breed with a single SNP mixed linear model including an additive polygenic effect with a covariance described by a genomic relationship matrix. The model was implemented via the EMMAX software (Kang et al. 2010).

### Statistical model

Based on their IDs, SNPs were annotated to genes corresponding to the ARS-UCD1.2 reference genome using Bioconductor BioMart tool version 3.14 (Smedley et al. 2009) and then genes were annotated to KEGG reference pathways (map) using the David software version 6.8 (Huang et al. 2007). The effects of KEGG pathways on stature were estimated separately for each breed using the following mixed linear model that accounted for the similarity between pathways:

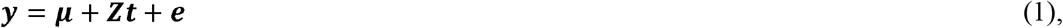

where *y* is the vector of absolute values of SNP additive effects on stature estimated in GWAS of Bouwman et al. (2018), *μ* represents the general mean, *t* is the random effect of KEGG pathways with a preimposed normal distribution defined by 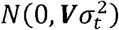 is a vector of residuals distributed as 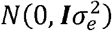, *Z* is an incidence matrix for *t*. Note that if multiple SNPs were identified within a gene only one SNP with the highest effect was included in *y*, so that each gene is represented by a single variant. The similarity between KEGGs *i* and *j*, was introduced into the model by incorporating a nondiagonal KEGG covariance matrix *v*. This covariance was expressed by the Jaccard similarity coefficient:

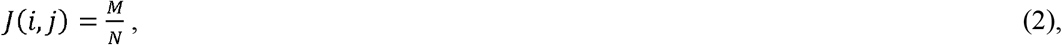

where *M* represents the number of genes shared between KEGG *i* and *j*, while N represents the total number of genes involved in KEGG *i* and *j*. Variance components were assumed as known, amounting 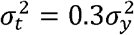 and 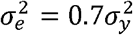.

### Solutions

The mixed model equations (Henderson 1984) were used to obtain solutions for *μ* and *t*:

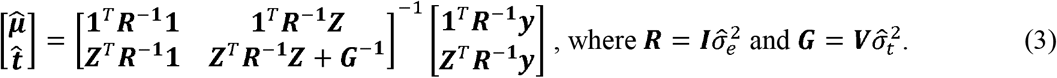

To maximise the computational performance of the estimation/prediction process, a custom Python program implementing the NumPy 1.19.5 library (Harris et al. 2020) was used. Since all calculations were carried out on a high-performance server, the NumPy library was also used to set the array indexing and order which further improved the computing time compared to a native Python application. Each element of 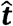 was assessed for significance 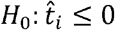 vs.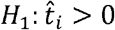 by calculating the probability of obtaining a more extreme value from the 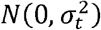 density function.

Since NumPy and SciPy APIs are implemented with LAPACK and BLAS, which require Fortran memory layout, all input matrices were transformed to Fortran order to avoid costly transposing. In comparison to a fixed matrix input, this approach results in a ten times faster estimation process.

## Data Access

Accession codes are available at https://doi.org/10.1038/s41588-018-0056-5.

## Acknowledgment

The genome-wide association studies were part of the Center for Genomic Selection in Animals and Plants (GenSAP) financed by Innovation Fund Denmark (Grant: 0603-00519B). The 1000 Bull Genomes Project is acknowledged for sharing sequence data for imputation. We thank prof. Bernt Guldbrandtsen for fruitful discussions on pathway modelling.

Calculations have been carried out using resources provided by Wroclaw Centre for Networking and Supercomputing.

